# Simple, Fast and Highly Efficient One-or Two-step Proteomic Preparation Enables Deep Profiling of Microgram-level FF and FFPE Tissues

**DOI:** 10.1101/2025.10.09.678953

**Authors:** Chuping Wei, Qiuxia Zhang, Changying Fu, Yeye Leng, Chuanxi Huang, Fuchu He, Yun Yang

## Abstract

Large-scale tissue proteomics requires workflows that are efficient, rapid, and repeatable across diverse samples. Herein, we present Simple Workflow for Integrated and Fast Tissue-preparation (SWIFT), which enables complete processing of fresh-frozen (FF) and formalin-fixed, paraffin-embedded (FFPE) tissues in either one- or two-step formats, while maintaining deep proteome coverage with high repeatability from low to microgram-level tissues. For FF tissues, an incubation process integrating lysis, reduction, alkylation, and digestion generates peptide samples directly from tissues in ≤ 1.5 h. For FFPE tissues, concurrent deparaffinization, rehydration, and de- crosslinking is achieved within 0.5 h, followed by one-step peptide preparation. Furthermore, our workflows eliminate desalting and offline cleanup steps, thereby reducing variability and total processing time. Using our methods, we identified up to ∼10,000 protein groups and ∼150,000 peptides across multiple mouse organs on timsTOF Pro. Repeatability was high (pairwise Pearson’s r > 0.96 across six experimental replicates), with dynamic ranges spanning 6–7 orders of magnitude. Organ-enriched protein analysis identified functionally distinct proteins unique to each tissue Paired FF and FFPE analyses revealed preservation-induced shifts, with FFPE tissues showing reduced detection of membrane-associated and respiratory proteins, including mitochondrial Complex I. Together, our fast and simplified workflows enable deep tissue proteomics for large-scale clinical and translational studies in a cost-effective and widely-accessible manner.

## INTRODUCTION

Mass-spectrometry–based tissue proteomics provides deep, quantitative profiling across clinical cohorts, enabling discovery and validation of protein biomarkers, pathway-level analyses of pathophysiology, and network-level mapping of disease mechanisms^1–4^. Fresh-frozen (FF) tissue preserves proteins in a state close to native conformation, enabling high-fidelity proteome measurements. By contrast, formalin-fixed, paraffin-embedded (FFPE) tissue offers low-cost, room-temperature storage while maintaining histoarchitecture for pathology and retrospective studies. Because both FF and FFPE formats are widely used in clinical research, extracting maximum value from them requires pipelines that are fast, robust, and highly sensitive.

Over the past two decades, advances in tissue sample preparation and mass spectrometric instrumentation have expanded tissue proteomics, enabling deep and repeatable profiling across large clinical cohorts. Several classical workflows were initially developed for cell samples but were later extended and adapted for tissue processing. Filter-Aided Sample Preparation (FASP) and Single-Pot, Solid-Phase-Enhanced Sample Preparation (SP3) are filter/bead-based buffer-exchange protocols^5,6^. These methods retained or immobilized proteins or peptides to remove detergents and chaotropes while restoring liquid chromatography–mass spectrometry (LC–MS) compatibility, but they typically required repeated binding–wash–elution cycles. In-StageTip (iST) and simple and integrated spintip-based proteomics technology (SISPROT) further streamlined the workflow by integrating reduction, alkylation, digestion, cleanup, and fractionation steps within a single tip^7,8^. For biopsy-level tissue samples, new sonication-based technologies have been developed, such as Pressure Cycling Technology (PCT) and Heat ‘n Beat (HnB)^9,10^. Specifically, PCT utilized alternating cycles of ultrahigh and low hydrostatic pressure in a miniaturized, disposable mechanical tissue homogenizer (MicroPestle) to facilitate rapid protein extraction. In a mouse multi-organ study, a PCT–Sequential Window Acquisition of All Theoretical Mass Spectra (PCT–SWATH) pipeline quantified on average 7,472 proteins in spleen, 6,911 in kidney, and 6,064 in lung on a Q Exactive HF-X^11^. HnB incorporated heating to reduce protease activity and bead-beating to improve homogenization efficiency, optimizing conditions for protein extraction and enabling an efficient end-to-end process. In a human cancer biopsy cohort, ∼4000−6000 protein groups were quantified. Sample Preparation by Easy Extraction and Digestion (SPEED) employs neat trifluoroacetic acid (TFA) for rapid lysis/inactivation, followed by neutralization with Tris base; upon neutralization, proteins form fine suspended particulates, allowing reduction, alkylation, and tryptic digestion in the same tube without solid-phase cleanup^12^. Although existing methods have been proven valuable in clinical research, their numerous handling steps, long processing time, and high costs hinder scalability for large cohorts and wider applications. Most existing approaches still rely on contaminant removal strategies to ensure cleaner downstream LC-MS analysis. However, peptide purification steps—including desalting, vacuum drying, and resuspension—remain manual and time-consuming. Therefore, simplified, robust, and repeatable workflows are urgently needed to facilitate sample preparation, shorten processing time, increase throughput, and improve reliability, enabling broader and more convenient clinical applications.

FFPE tissue proteomics requires additional deparaffinization and retrieval of cross- linked proteins prior to sample preparation. A highly efficient de-crosslinking process is crucial for downstream protein solubilization and enzymatic digestion^13^. To overcome extraction barriers caused by crosslinking, several streamlined workflows have been developed. In an FFPE-adapted FASP workflow, deparaffinized tissue was lysed in sodium dodecyl sulfate (SDS)/dithiothreitol (DTT) at 99 °C for 1 h, followed by on-filter SDS-to-urea exchange for cleanup^14^. Using simple pipette-tip fractionation, this workflow identified 5,203 protein groups from FFPE mouse liver in ∼ 24 h. Yi Zhu et al. developed an FFPE-compatible PCT–SWATH workflow^15^: FFPE tissue sections underwent xylene dewaxing and graded-ethanol rehydration, followed by sequential acidic and basic hydrolysis, and pressure-cycling–enhanced extraction and digestion. Meanwhile, xylene-free and simplified workflows were developed to accelerate pretreatment. The HnB method accelerated dewaxing and decrosslinking by combining brief heptane–methanol deparaffinization in Barocycler tubes (≤ 12 min) with controlled heating and bead-beating^10^. Georgia Mitsa et al. employed hot-water deparaffinization to dissolve paraffin without toxic solvents, achieving protein yields and stability comparable to xylene-based methods^16^. Magdalena Kuras et al. combined xylene-free paraffin removal with antigen retrieval at 97 °C, and optimized two-enzyme digestion with suspension trapping (S-Trap)^17^. High-yield protein extraction and recovery by direct solubilization (HYPERsol) integrated direct SDS extraction, adaptive focused acoustics (AFA) sonication, and S-Trap on-trap digestion (∼1 h) to enable thermal retrieval/decrosslinking and rapid peptide generation^18^. However, these workflows rely on multiple handling steps and transfers, which prolong processing time, introduce variability, reduce sensitivity, and limit scalability. Moreover, many of these methods require specialized devices and consumables, increasing cost and restricting accessibility.

Despite advances, tissue proteomics remains constrained by multi-step operations and transfers, which might limit throughput, stability and sensitivity. Herein, we introduce two highly efficient workflows—One-step SWIFT (Simple Workflow for Integrated and Fast Tissue-preparation) for FF tissues and Two-step SWIFT for FFPE tissues. For FF tissues, we integrated all the sample preparation steps into a single incubation step to realize lysis, reduction, alkylation, and digestion simultaneously. For FFPE tissues, an extra one-step pretreatment achieves near-complete deparaffinization, rehydration, and decrosslinking. By integrating all the necessary steps into highly simplified workflows, SWIFT workflows minimize handling complexity and sample loss while demonstrating high sensitivity, high repeatability, and deep proteome coverage. Total processing times are merely ∼1.5 h for FF tissues and ∼2 h for FFPE tissues, and the resulting samples are ready for direct liquid chromatography–tandem mass spectrometry (LC-MS/MS) analysis. Using our SWIFT workflows, we systematically evaluated proteome performance and fixation-linked bias in paired FF and FFPE tissues across seven mouse organs. Together, our SWIFT workflows aim to lay a foundation for large-scale proteomic studies across diverse tissue types and preservation modalities.

## EXPERIMENTAL SECTION

### One-step SWIFT Workflow for FF tissues

The FF tissues were washed three times with 1 ×phosphate-buffered saline (PBS) at 4 °C to remove contaminants. Next, 30 µL of a mixed-component buffer was added to each tissue fragment, and 10 µL to slide-mounted tissue sections. The mixed-component buffer contained rapid trypsin/Lys-C (0.335 µg/µL), tris(2-carboxyethyl) phosphine hydrochloride (TCEP-HCl, 1 mM), 2- Chloroacetamide (CAA, 4 mM), and n-Dodecyl-β-D-maltoside (DDM, 10%) in rapid digestion buffer. Samples were sonicated for 10 min (20 s on/20 s off, 85% amplitude) using a Qsonica Q800R3 sonicator until fully homogenized, followed by incubation at 70 °C for 1 h. Note, an additional boiling step at 95 °C was applied to pancreatic tissues to denature endogenous digestive enzymes and to prevent autolysis. Peptide concentration was measured at 205 nm using a NanoDrop One spectrophotometer (Thermo Scientific). Samples were diluted to 100 ng/µL in 0.1% FA for nanoLC– MS/MS analysis.

### Two-step SWIFT Workflow for FFPE tissues

For pretreatment, two slides of FFPE tissue were incubated with 0.5 mL of 300 mM Tris(hydroxymethyl)aminomethane (Tris)–HCl buffer (pH 8.5) at 95 °C for 10 min with shaking (1500 rpm) on a thermomixer. During this step, molten paraffin formed a turbid suspension with the aqueous buffer, while the tissue fragments underwent rehydration and heat- and base-induced de-crosslinking. After incubation, samples were centrifuged at 4 °C for 10 min. A distinct paraffin layer floated on the surface of the aqueous phase, while the tissue fragments pelleted at the bottom. Tissue pellets were transferred into new tubes for subsequent lysis and peptide preparation. The following process was identical to that described above in the “’One-step SWIFT Workflow for FF Tissues’” section, except that for slide-derived tissue samples, peptides were passed through a cell-strainer to remove residual tissue fragments.

### Conventional multi-step pretreatment for FFPE samples

FFPE sections were briefly centrifuged to the bottom of the tubes. Deparaffinization solution (500 µL) was added, followed by 10-min incubation at 37°C and 800 rpm, and subsequent centrifugation at 16,000 × g for 3 min. Next, the supernatant was discarded. This deparaffinization step was repeated 2–3 times. A graded series of ethanol solutions (100%, 90%, and 75%) were sequentially applied (500 µL each). Each rehydration step included 5-min incubation at 37°C and 800 rpm, followed by 3-min centrifugation at 17,000 × g and supernatant removal. Samples were then rehydrated with two water rinses (500 µL each) at 37 °C, 800 rpm, for 2 min, followed hy centrifugation at 17,000 ×g for 5 min. After the final rehydration step, samples were air-dried for 5 min in open tubes at 65 °C. Finally, decrosslinking buffer (300 mM Tris-HCl, 1% sodium deoxycholate (SDC), pH 8.8) was added at 10 volumes relative to the tissue, and samples were incubated at 95°C, 400 rpm, for 90 min, followed by brief centrifugation to collect condensates.

### Conventional multi-step sample preparation workflow

Following FFPE tissue pretreatment, proteins were extracted by sonication (Qsonica Q800R3) at 85% amplitude (4 s on/10 s off cycles) for 3 min. Then, samples were cooled, and protein concentration was quantified by bicinchoninic acid assay (BCA) method. Reduction and alkylation were performed by adding TCEP to a final concentration of 10 mM and CAA to 40 mM, followed by incubation at 65 °C for 15 min. For enzymatic digestion, MS-grade trypsin (in 100 mM Tris–HCl, pH 8.8) was added at an enzyme-to-protein ratio of 1:33 (w/w), and samples were incubated at 37 °C overnight. After digestion, peptide solutions were acidified to ∼pH 2 with FA. The supernatant was collected after centrifugation. Peptides were then desalted using C18 reversed-phase solid-phase extraction procedure. Eluates were dried by vacuum centrifugation. Peptide concentration was determined by absorbance at 205 nm (NanoDrop One), and samples were reconstituted to 100 ng/µL with 0.1% FA for nanoLC–MS/MS analysis.

## RESULTS AND DISCUSSION

### Development of One-step SWIFT Workflow for FF Tissues

Multicomponent Reactions (MCRs) in organic chemistry are one-pot synthetic processes where three or more reactants are combined simultaneously to synthesize a single product^19,20^. The popularity of MCRs lies in their simplicity and high efficiency in resource utilization (e.g., time, energy, and materials). Inspired by the concept of MCRs, we aim to adapt this strategy to the sample preparation process for bottom-up proteomics. Traditionally, biological samples undergo sequential steps before LC-MS analysis—such as cell lysis, protein extraction, disulfide bond reduction, alkylation, enzymatic digestion, desalting, vacuum dryness, and reconstitution. Each subsequent step typically begins only after the completion of reactions in the preceding step and the removal or quenching of excess reagents, which makes the overall multi-step process time-consuming and labour-intensive (Fig.S1a). Meanwhile, multiple sample transfers and stepwise reagent additions could increase the risk of sample contamination. By leveraging the MCR concept, we propose integrating these traditionally sequential procedures into a single integrated system. Our approach combines all necessary reagents in a single vessel and eliminates all unnecessary procedures, allowing them to react concurrently, thereby reducing manual handling and technical variability, and improving throughput (Fig. 1a, Fig.S1a).

**Figure 1.**
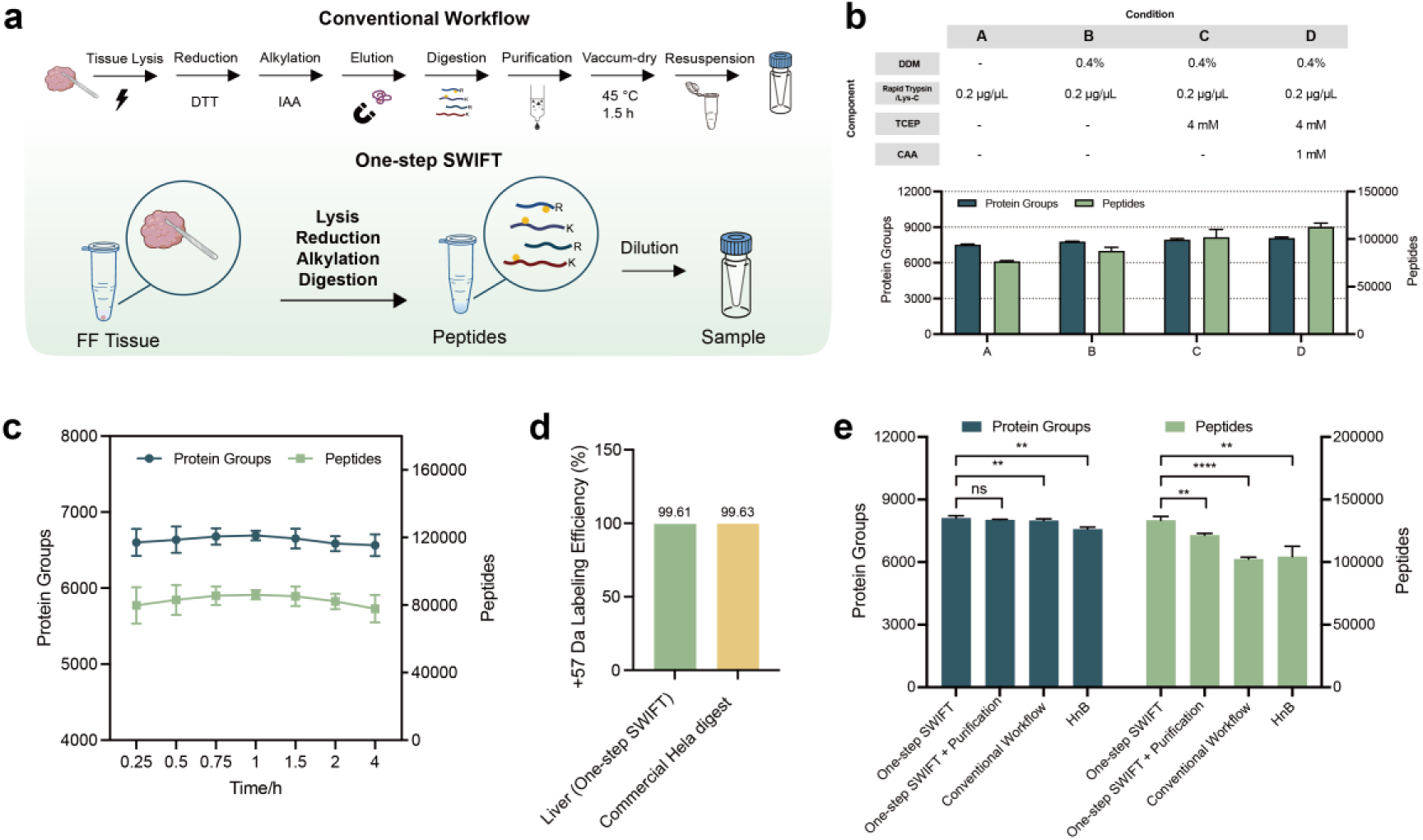
Development of the One-step SWIFT workflow for FF tissues. (a) Scheme of the One-step SWIFT sample preparation workflow for FF tissues, compared with a conventional multi-step workflow. Multicomponent buffer lyses, reduces, alkylates and digests FF tissues into peptides in a single step. After simple dilution, samples are ready for nanoLC–MS/MS. (b) Optimization of the multicomponent buffer composition. (c) Optimization of incubation times. (d) Comparison of cysteine alkylation efficiency between liver samples processed by One-step SWIFT and a commercial HeLa digest. (e) End-to-end benchmarking of One-step SWIFT, One-step SWIFT with extra purification, and a conventional multi-step workflow. Statistical differences were assessed by two-sided Student’s t-test.

This one-step workflow enables direct and fast tissue-to-peptide preparation. After rinsing in PBS, a mixed-component buffer is added to the tissue, followed by sonication to homogenize the tissue and to extract proteins. The resulting peptide samples are ready for nanoLC–MS analysis. The entire process requires only a single addition of the mixed-component buffer, minimizing contamination risk and enhancing repeatability. Moreover, no specialized instrumentation, consumables or reagents are required, making the workflow economical and widely accessible.

We first optimized mixed-component buffer composition to establish an efficient and effective tissue-to-peptide workflow. Around 300 µg wet mouse liver tissue was used for optimization per condition (n = 3). The mild, sugar-based detergent DDM was included to enhance protein solubilization, while rapid trypsin/Lys-C was used to accelerate proteolysis. In addition, CAA and TCEP were used for reduction and alkylation. Our results demonstrated that the composition consisting of all the above four reagents yielded the highest numbers of proteins and peptides (Fig. 1b). In addition, rapid trypsin/Lys-C outperformed conventional MS-grade trypsin at their respective optimal temperatures (70 °C vs 37 °C), and an enzyme concentration at 0.033 μg/μL maximized protein numbers while minimizing enzyme use (Fig. S1b).

To evaluate the reactivity of the integrated process, seven different time points (0.25, 0.5, 0.75, 1, 1.5, 2, 4 h) were compared. Protein and peptide identifications reached their maximum at 1 h and exhibited the least variation, indicating the highest efficiency and stability (Fig. 1c). Miss-cleavage fluctuated around 38 % and followed the same trend (Fig. S1c). By evaluating the alkylating efficiency, we observed that liver samples processed using our workflow showed comparable ratios of modified peptides to a commercial HeLa digest, confirming that our one-step method achieves similarly high alkylation efficiency (Fig. 1d). Subsequently, we compared four workflows: One-step SWIFT, One-step SWIFT with extra purification steps, a conventional multi-step workflow, and the recently published HnB method. Protein identifications showed no significant differences across the first three workflows, while One-step SWIFT achieved the highest number of peptides (Fig. 1e). Total-ion chromatograms revealed that residual reagent peaks remained in One-step SWIFT; however, peptide separation remained unaffected. Importantly, omitting desalting preserved more hydrophilic peptides, whereas C18 cleanup removed some of these peptides and could skew downstream quantification (Fig. S1d). Therefore, our One-step SWIFT integrates reactions within an hour with high efficiency and completeness. By omitting purification steps, it minimizes sample loss while achieving protein and peptide coverage comparable to or better than that of conventional multi-step workflows, enabling faster, simpler, and more efficient sample preparation.

### Deep Proteome Coverage Across Seven FF Mouse Organs Using One-step SWIFT

To evaluate the cross-organ robustness and proteome depth of our One-step SWIFT workflow, we profiled seven mouse organs—brain, heart, lung, spleen, kidney, liver, and pancreas—each with six replicates. The organs are arranged in a simplified physiological order (neural → contractile → respiratory/immune → excretory → metabolic/digestive). Single-run LC-MS/MS identified an average of 8,995 protein groups in the brain, 7,522 in the heart, 10,122 in the lung, 9,747 in the spleen, 8,569 in the liver, 9,107 in the kidney, and 8,985 in the pancreas. Mean peptide identifications per sample ranged from 91,680 to 158,335 (Fig. 2a). To prevent self-proteolysis from endogenous enzymes, pancreatic samples underwent a heat pretreatment, which effectively suppressed autolysis according to our results (Fig.S1e).

**Figure 2.**
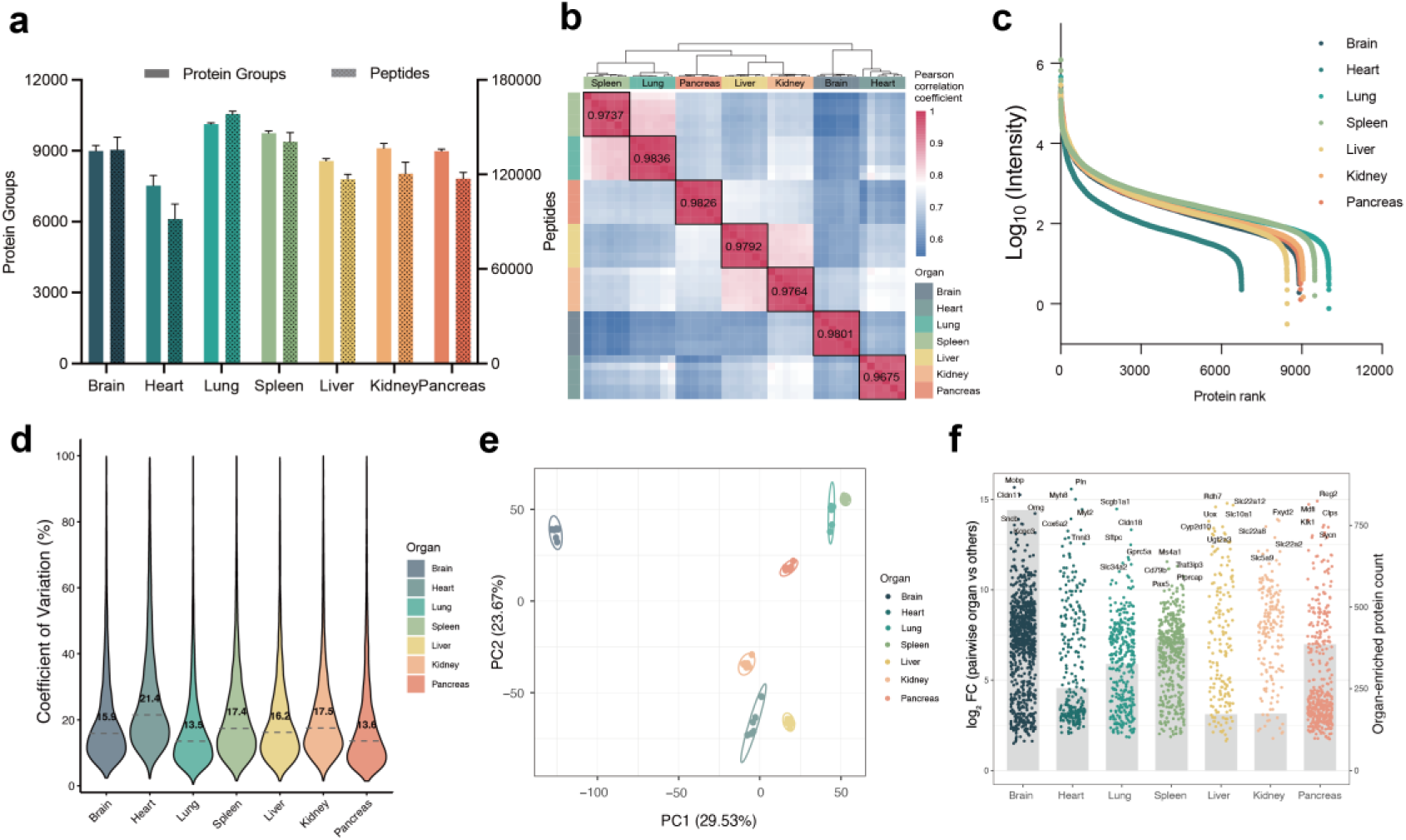
Performance of the One-step SWIFT workflow for FF tissues. (a) Protein group and peptide identifications for FF tissues using the One-step SWIFT workflow across seven mouse organs (n = 6). (b) Pearson correlation matrix with unsupervised clustering across organs. (c) Dynamic range of protein expression from seven FF tissues. (d) CVs across organs; medians are indicated in the violin plots. (e) PCA showing clear clustering among organs. (f) Volcano plots of pairwise organ-enriched proteins of FF tissues. Statistical testing used limma-treat (trend=TRUE) with thresholds FDR ≤ 0.01 and |log2FC| ≥ 1. The top five proteins by |log_2_FC| are labelled; organ-specific set sizes (protein counts) are shown on the plots.

The acquired proteomic data consistently demonstrated high repeatability, supporting the robustness of our One-step SWIFT workflow. Within-organ Pearson correlation coefficients were 0.974 (spleen), 0.984 (lung), 0.979 (liver), 0.976 (kidney), 0.980 (brain), 0.968 (heart), and 0.983 (pancreas) (Fig. 2b). Protein intensities spanned 6-7 orders of magnitude (Fig. 2c). Coefficients of variation (CVs) for protein groups, visualized as violin plots, showed median CVs ranging from 15.9% in lung and pancreas to 24.6% in heart, indicating low technical variability (Fig. 2d). Heart tissues showed a slightly lower number of identifications and higher CVs, likely due to residual blood contamination. Principal component analysis (PCA) effectively clustered replicates by organ, further highlighting the robustness of our workflow (Fig. 2e). The high repeatability and high quantitative precision in our data benefit from the design of our workflow, which reduces variance and contamination by minimizing sample transfers and operation steps.

To assess biological specificity in our dataset, we performed differential expression analyses across the seven organs. Organ-enriched protein expression is fundamental to organ biology, as each organ’s specialized function is underpinned by a distinctive proteome profile^21^. By comparing each organ against all others, we identified organ-enriched proteins as those selectively enriched in a given organ. The volcano plots summarize these comparisons (false discovery rate (FDR) < 0.01 and ≥2-fold change) and label the five proteins with the highest fold changes for each organ (Fig. 2f). Consistent with previous proteome-wide surveys, the brain exhibited the highest number of organ-enriched proteins, reflecting its complex and specialized cellular roles.

By contrast, organs such as the liver and kidney expressed far fewer uniquely expressed proteins–many of their key proteins (e.g., metabolic enzymes) are shared across multiple tissues. These results demonstrate that our method effectively captures biologically significant proteins without compromising proteome depth, despite its highly simplified workflow.

Gene Ontology (GO) biological process (BP) enrichment analysis further validated the biological relevance of these organ-enriched proteins (Fig. S2). In the brain, enrichment terms included neurotransmitter secretion, synaptic vesicle exocytosis, and glutamatergic/GABAergic transmission, consistent with neuronal function. Heart enrichment was dominated by oxidative phosphorylation, mitochondrial ATP synthesis, and related respiratory terms, aligning with high energetic demand. Lung enrichment included axoneme assembly, cilium-driven fluid movement, and endothelial/angiogenic programs, reflecting its ciliated epithelium and vascular network. The spleen showed enrichment for T-cell activation, T cell receptor (TCR) signalling, and B-cell regulation, consistent with its role in adaptive immunity. Liver-enriched terms—such as steroid/cholesterol biosynthesis, amino-acid and fatty-acid metabolism, monocarboxylate pathways—highlight hepatocyte-centered metabolic specialization. The kidney was enriched for renal absorption and broad ion/solute transport, consistent with tubular reabsorption and homeostasis. Pancreas-enriched terms included terms ribosome biogenesis and translational initiation/elongation, aligning with the heavy protein-secretory output of acinar cells.

Overall, these analyses confirm that our proteomic approach reliably identifies organ-enriched proteins and their associated biological processes, providing biologically credible insights with robust coverage and cost-effective simplicity.

### Development of Two-step SWIFT Workflow for FFPE tissues

We aimed to adapt the One-step SWIFT workflow for FFPE tissues while maintaining high efficiency and deep proteome coverage. Conventional FFPE pretreatment is labor-intensive, requires multiple tissue transfers, and impedes efficient peptide generation^22–24^. Moreover, such protocols rely on toxic and environmentally harmful reagents. To overcome these limitations, we developed a pretreatment method that simultaneously performs deparaffinization, rehydration, and de-crosslinking. In this pretreatment, a high-temperature alkaline buffer (95 °C) dissolves paraffin and reverses protein cross-links, followed by rapid centrifugation at 4 °C to separate tissues from the aqueous phase. This pretreatment is completed within 30 min, directly addressing time and manual handling bottlenecks. The overall preparation of FFPE tissues thus comprises a pretreatment step followed by One-step SWIFT, constituting the Two-step SWIFT workflow (Fig. 3a).

**Figure 3.**
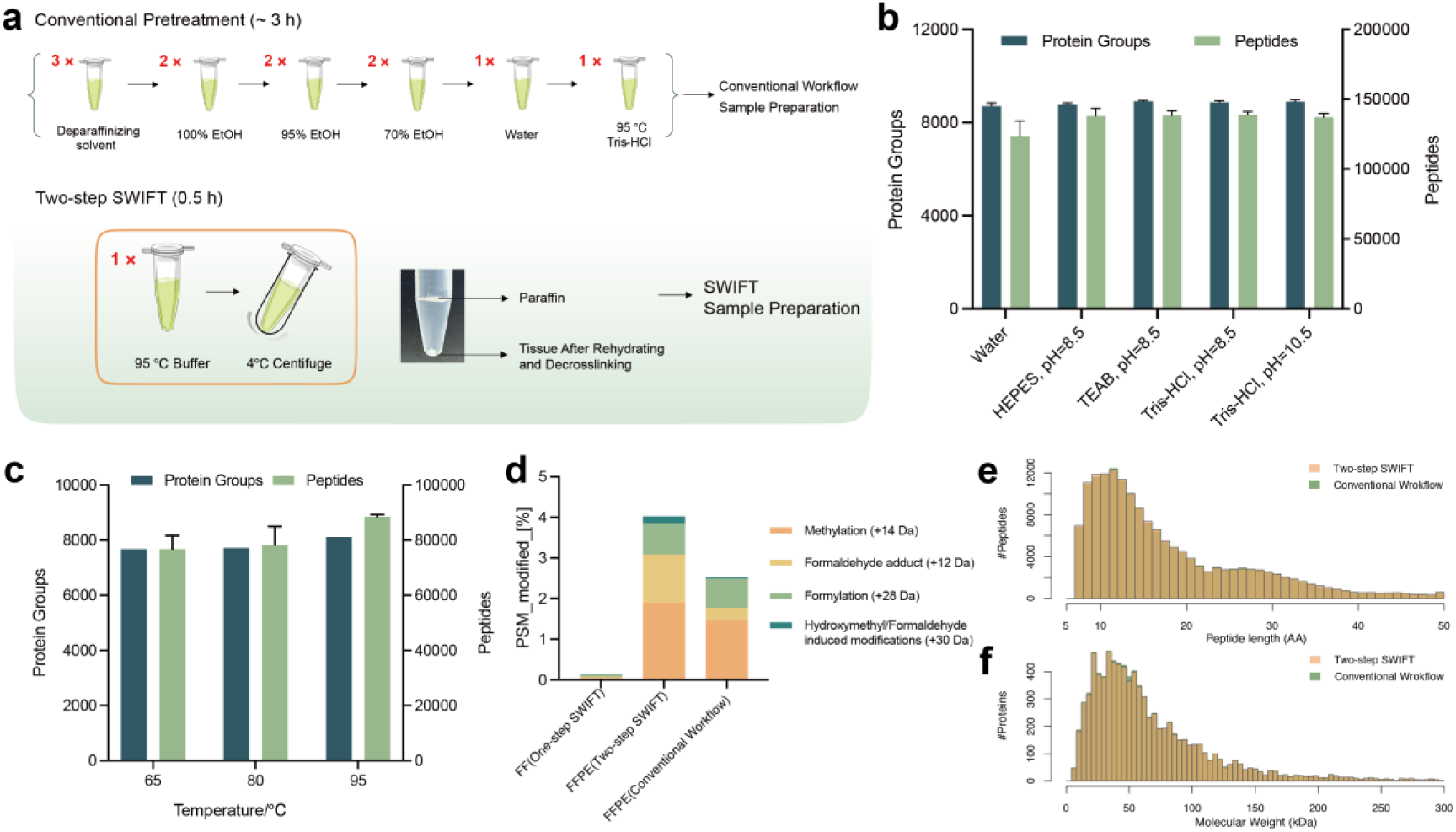
Development of the Two-step SWIFT workflow for FFPE tissues. (a) Comparison of the One-step SWIFT pretreatment for FFPE tissues to a conventional xylene-based multi-step pretreatment. (b) Protein-group and peptide identifications across buffer types and pH conditions. (c) Protein-group and peptide identifications across different incubation temperatures. (d) Percentages of fixation-induced modifications in FF (One-step SWIFT), FFPE (Two-step SWIFT), and FFPE (conventional multi-step workflow). (e) Comparison of cysteine alkylation efficiency in FF liver, FFPE brain, and a commercial HeLa digest.

The buffer composition and experimental conditions were systematically optimized to maximize performance. For optimization, two 4-µm FFPE mouse brain sections were used per condition (n = 3). We first compared several commonly used buffers across different pH values (Fig. 3b). Except for water, all alkaline buffers yielded comparable protein and peptide identifications, supporting the conclusion that alkaline conditions effectively reverse cross-linking. Additives such as SDC, hydroxylamine, and formaldehyde dehydrogenase (FDH) were also tested but had minimal impact on protein group or peptide identifications (Fig. S3a). We then independently evaluated incubation time and temperature, revealing that a single-step incubation at 95 °C for 10 min was sufficient to maximize both protein group and peptide identifications (Fig. 3c, S3b).

To verify the effectiveness of the pretreatment in reversing crosslinking, we systematically assessed fixation-induced modifications, peptide physicochemical properties, and alkylation efficiency in FFPE tissues. An open-search analysis of data independent acquisition (DDA) data with MSFragger identified four formaldehyde-induced mass shifts among the ten most abundant modifications: methylation (+14 Da), formaldehyde adduct (+12 Da), formylation (+28 Da), hydroxymethyl/formaldehyde-induced modifications (+30 Da) (Fig. S3d)^25,26^. These formaldehyde-induced peptide-spectrum matches (PSMs) accounted for less than 4% of spectra, representing a minimal modification effect, although the conventional multi-step workflow can further reduce these PSMs (Fig. 3d). The observed rate of crosslinking-related modifications was consistent with published reports, indicating that our simplified workflow maintains comparable performance^27^. Carbamidomethylation (+57 Da) in FFPE brain was only slightly lower than that observed in a commercial HeLa digest, yet remained high, confirming robust alkylation efficiency (Fig. S3c). Fig. 3e and 3f show similar molecular weight and peptide length distributions between the Two-step SWIFT and the conventional multi-step workflow, indicating their overall comparability.

In summary, we integrated multiple manual deparaffinization, rehydration, and de- crosslinking steps into a single incubation, achieving a total processing time of roughly 30 min and confirming effective cross-link reversal by open-search modification analysis.

### Deep Proteome Coverage Across Seven FFPE Mouse Organs Using Two-step SWIFT

To evaluate cross-organ robustness and proteome coverage of Two-step SWIFT workflow, we analyzed the same seven mouse organs used throughout this study. Single-run LC-MS/MS (n = 6 per organ) identified, on average, 8,960 protein groups in the brain, 6,961 in the heart, 9,027 in the lung, 8,866 in the spleen, 8,156 in the liver, 9,131 in the kidney, and 8,023 in the pancreas. Across organs, the mean peptide identifications per sample ranged from 79,663 to 134,848 (Fig. 4a).

**Figure 4.**
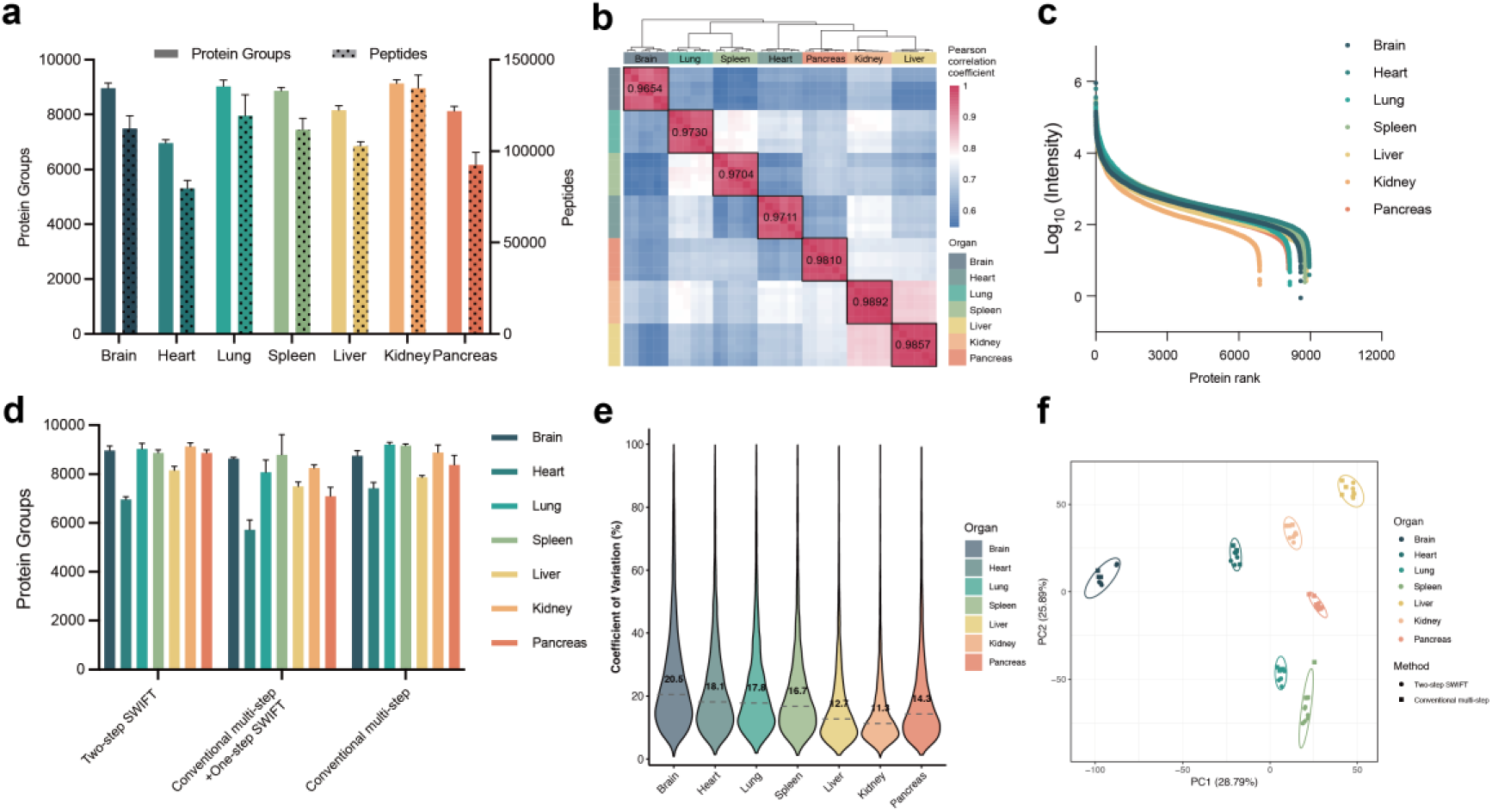
Performance of the Two-step SWIFT workflow for FFPE tissues. (a) Protein group and peptide identifications for FFPE tissues using the Two-step SWIFT workflow across seven mouse organs (n = 6). (b) Pearson correlation matrix with unsupervised clustering across organs (c) Dynamic range of protein expression from seven FFPE tissues. (d) Benchmarking FFPE tissue workflows for proteome coverage. Protein group counts for each organ are shown for: Two-step SWIFT (n = 6), conventional multi-step pretreatment plus One-step SWIFT (n = 3), and conventional multi-step pretreatment plus conventional multi-step sample preparation (n = 3). (e) Percentages of CVs across organs; medians are showed in the violin plots. (f) PCA showing clear clustering among organs regardless of different workflows.

The acquired proteomic data consistently demonstrated a high degree of repeatability, supporting the robustness of the Two-step SWIFT workflow. Within-organ Pearson correlation coefficients were 0.965 (brain), 0.971 (heart), 0.973 (lung), 0.970 (spleen), 0.986 (liver), 0.989 (kidney), and 0.981 (pancreas) (Fig. 4b). Protein intensities spanned 6-7 orders of magnitude, demonstrating the capability to detect low-abundance proteins (Fig. 4c). This dynamic range was comparable to that achieved in FF tissues from the previous analysis (Fig. 2c).

To benchmark processing time and procedural complexity, we compared three FFPE workflows: (i) Two-step SWIFT (n = 6); (ii) conventional multi-step pretreatment plus One-step SWIFT (n = 3); and (iii) conventional multi-step pretreatment plus conventional multi-step sample preparation (n = 3). Across most organs, Two-step SWIFT delivered deeper proteome coverage than conventional pretreatment plus One-step SWIFT and achieved protein-group identifications comparable to the conventional multi-step workflow (Fig. 4d). CVs for protein groups ranged from 11.3% (kidney) to 20.5% (brain), indicating low technical variability (Fig. 4e). CVs obtained using the conventional multi-step workflow were similar to those for Two-step SWIFT, indicating comparable technical precision (Fig. S4a). PCA grouped replicates by organ regardless of workflow, underscoring the repeatability of Two-step SWIFT and its comparability to the conventional multi-step workflow (Fig. 4f).

Furthermore, to evaluate biological coherence, we repeated the cross-organ contrasts in the FFPE samples using the same pairwise framework (FDR < 0.01; ≥2-fold). We again identified organ-enriched protein sets, whose sizes and distributions closely mirrored those in the FF data (Fig. S4b). This concordance indicates that the organ- defining proteomic signals are preserved across tissue preservation methods. GOBP enrichment of the organ-enriched sets converged on canonical programs for each tissue and remained almost consistent across FF and FFPE datasets (Fig. S4c). Brain terms were enriched for synaptic transmission and neurotransmitter secretion; heart for oxidative phosphorylation and ATP synthesis; lung for cilium/axoneme assembly and angiogenesis; spleen for T-cell activation and B-cell regulation; liver for lipid/cholesterol and amino-acid metabolism; kidney for ion/solute transport and renal absorption; pancreas for ribosome biogenesis/translation, reflecting its high secretory output. The recurrence of these top-ranked terms—along with large intersection sizes and strong −log10(q) values—indicates that the organ-enriched protein signatures reflect true biological patterns. Overall, the Two-step SWIFT workflow identified organ-enriched proteins and pathways with deep coverage and high repeatability across preservation methods, while remaining operationally simple and scalable.

### Evaluating Preservation Effects on the Mouse Proteome

Despite the clinical ubiquity of FFPE tissues, fixation and embedding procedures alter the measured proteome compared to fresh-frozen tissues. During fixation, formalin permeates cellular compartments and stabilizes proteins by forming crosslinks, classically methylene bridges between proximal side chains, primarily at lysine residues^28^. While this enables stable long-term preservation, the fixation chemistry alter the proteome at multiple levels: (i) at the level of primary and secondary protein structure, where sequence modifications impair database-search recognition — bioinformatics analyses suggest that ∼6–10% of peptides are modified ^27,25^, and (ii) at the level of higher-order structure, where cross-linking disrupts α-helices and β-sheets, changes tertiary conformation, and promotes quaternary aggregation^29,30^. Together, these effects reduce solubility and increase resistance to conventional extraction buffers, thereby constraining downstream sample processing and protein identification. Previous studies reported ∼91–92% overlap in protein identifications between FFPE and FF proteomes^14,31^, with Pearson correlation coefficients of ∼0.91-0.93^14,17^. These findings demonstrate preservation-induced bias and underscore the need to systematically measure and account for fixation effects to interpret biological signals accurately.

We first compared proteome depth and global proteome similarity between paired FF and FFPE tissues across seven organs. Using adjacent halves of the same organs, we applied One-step and Two-step SWIFT workflows to enable a controlled comparison across preservation methods. We identified 11,957 protein groups (1% FDR), corresponding to 11,887 protein-coding genes in the mouse UniProt Reference Proteome^32^. Overall, FF tissues yielded 4–9% more protein identifications than FFPE: brain (+8.53%), heart (+8.69%), lung (+7.68%), spleen (+4.18%), liver (+4.58%), kidney (+4.81%), and pancreas (+5.10%) (Fig. 5a). Pearson correlation coefficients between paired FF and FFPE tissues ranged from 0.859 to 0.917 across organs, modest values given the deep coverage achieved (Fig. 5b). These values are consistent with previous reports using multi-step workflows, underscoring the robustness of the measurements across preservation methods^14,17^. While PCA revealed clustering primarily by organ, preservation method also had a clear effect, indicating substantial fixation-associated shifts (Fig. 5c).

**Figure 5.**
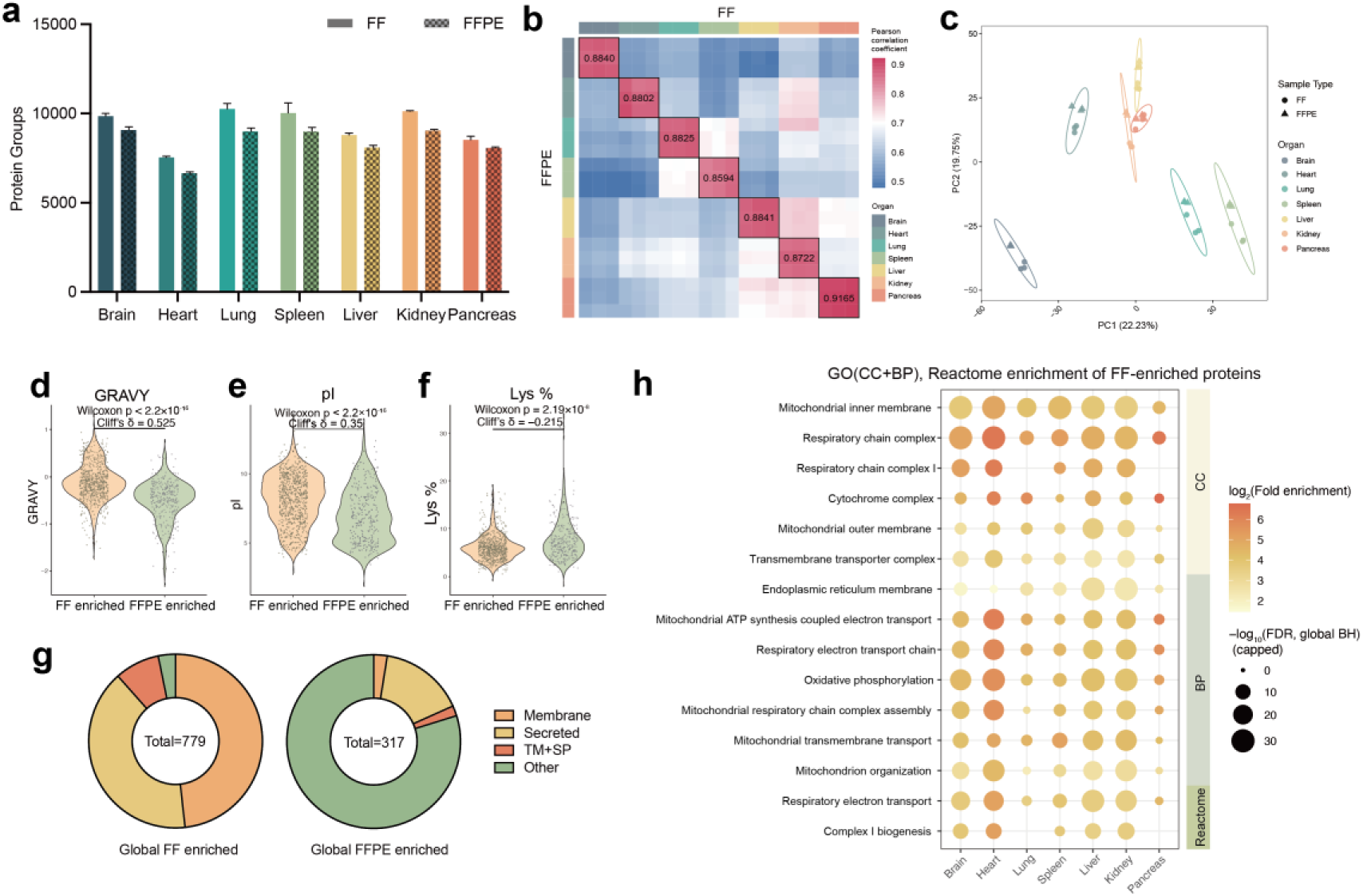
Comparison of paired FF and FFPE tissues to evaluate fixation and crosslinking effects. (a) Protein groups identified across paired FF and FFPE tissues (n = 3). (b) Pearson correlations among FF and FFPE tissues. (c) PCA analysis of paired FF and FFPE tissues across organs. (d–f) Distributions of pI (d), GRAVY (e), and Lys % (f) for FF-enriched and FFPE-enriched proteins. Statistical significance was determined using two-sided Wilcoxon rank-sum test with Cliff’s delta effect sizes. (g) Composition of enriched proteins by subcellular class (Membrane, Secreted, TM+SP, Other), based on consensus predictions. (h) GOBP, GOCC and Reactome over-representation analysis for FF-enriched proteins by organ.

We next examined the preservation effects reflected by FF–FFPE correlations and the clear FF–FFPE separation observed in PCA. First, we fit a global linear model controlling for organ-enriched effects to identify proteins consistently influenced by preservation methods. Proteins with |log₂FC| ≥ 0.8 and adjusted p-value ≤ 0.05 (Benjamini–Hochberg (BH) correction) were considered differentially abundant. Under this criterion, 779 protein groups were enriched in FF tissues and 317 in FFPE, indicating a pervasive but bidirectional preservation effect—aligned with the modest 4–9% advantage in FF identifications. To assess organ-enriched effects, we then performed within-organ contrasts with a stricter threshold (|log₂FC| ≥ 1.0, adjusted p-value ≤ 0.01) and visualized results in multi-organ volcano plots (Fig. S5a). This framework allowed us to distinguish global and organ-level differences, facilitating subsequent investigation of their sources and biological impact. To investigate whether crosslinking preferentially affects proteins based on structural or physicochemical features, we analyzed the differentially abundant global protein sets. We first compared distributions of hydropathy value (GRAVY), isoelectric point (pI), and lysine (Lys) content (%) using violin plots (Fig. 5d–5f). Compared to the FF-enriched set, FFPE- enriched proteins exhibited significantly lower GRAVY scores, lower pI values, and reduced Lys content. These shifts indicate that fresh-frozen storage better preserves hydrophobic proteins, consistent with previous findings^33^. We assume that the GRAVY shift reflects differences in membrane association. Using consensus annotations from three predictors (DeepTMHMM^34^, TOPCONS^35^, and SCAMPI^36^) and SignalP6.0^37^ for secreted-protein prediction, we applied a majority-decision method to assign membrane-protein topology. Over 56.6% of FF-enriched proteins were annotated as membrane-associated or containing both transmembrane and signal peptide domains. In contrast, 79.8% of FFPE-enriched proteins fell into other categories, consistent with soluble proteins; and an additional 15.77% predicted as secreted proteins (Fig. 5g). Collectively, these findings indicate a preservation-dependent bias: FFPE favors more acidic, and soluble proteins, whereas FF favors hydrophobic, membrane-associated proteins—consistent with known crosslinking and extraction mechanisms.

To further examine these biases, we performed per-organ over-representation analysis (ORA; hypergeometric test with Benjamini–Hochberg FDR) of FF- and FFPE- enriched protein sets and summarized non-redundant GOBP, GO cellular component (CC), and Reactome terms with effect sizes (log₂ fold-enrichment, log₂FE). Across multiple organs, FF-enriched proteins consistently mapped to mitochondrial membrane structures and energy production pathways, particularly oxidative phosphorylation (OXPHOS) and electron transport (Fig. 5h). Despite organ-enriched differences in enrichment magnitude, terms such as mitochondrial inner membrane, respiratory chain complex, and Complex I biogenesis remain consistently overrepresented. This pattern reflects the preferential detection of large, hydrophobic, multi-subunit complexes embedded in mitochondrial membranes—structures that are poorly recovered from FFPE tissues due to crosslink-induced rigidity and reduced solubility. Enrichment of transmembrane complexes, including the mitochondrial carrier MPC2, further supports the membrane-centric nature of the FF-enriched proteome (Fig. S5a). In contrast, FFPE-enriched terms clustered around soluble metabolic and stress-response pathways—oxidative/chemical stress, aldehyde and nicotinamide-nucleotide metabolism, amino-acid and carboxylic-acid catabolism (Fig. S5b). This aligns with known extraction biases: while formalin crosslinks can often be reversed in soluble proteins, they hinder digestion and recovery of hydrophobic membrane assemblies. Together, these data suggest that FFPE is more effective for recovering soluble and secreted enzyme networks, while FF excels at capturing hydrophobic, multi-pass membrane proteins—especially membrane-embedded OXPHOS machinery.

Our findings demonstrate that comparison of paired FF and FFPE tissues enables the detection and quantification of preservation-induced biases at scale. Formaldehyde fixation introduces methylene bridge crosslinks, primarily through lysine side chains, together with additional adducts that impede protein solubilization and limit protease accessibility. Subsequent paraffin embedding and dehydration further consolidate these crosslinked networks, creating formidable barriers to efficient protein extraction and digestion. The implications are clear: although FFPE tissues are invaluable for retrospective clinical and translational research, they carry a systematic underrepresentation of membrane systems, especially mitochondrial OXPHOS assemblies. Such biases need to be explicitly considered in data interpretation to avoid overlooking mechanistically important protein classes.

## CONCLUSIONS

Here we presented the simple SWIFT workflows for FF and FFPE tissues, enabling deep proteome coverage. Lysis, reduction, alkylation, and digestion were integrated into a single step for peptide preparation, while conventional “elution–drying– resuspension” procedures were omitted. For FFPE tissues, we simplified the deparaffinization, rehydration, and decrosslinking pretreatment processes into a single step. Our One-step and Two-step SWIFT workflows enable fast and high efficiency proteomic sample preparation with reduced variation and sample loss. From tissues to peptide samples, the total processing time was less than 1.5 h for FF tissues and less than 2 h for FFPE tissues. Using the SWIFT workflows, deep proteome coverage was achieved across seven mouse organs using a timsTOF Pro: 7,535–10,253 protein groups per FF tissue and 6,658–9,082 per FFPE tissue. Comparative analysis of paired FF and FFPE tissues revealed consistent enrichment of respiratory chain and OXPHOS-related terms in FF samples, indicating that fixation chemistry in FFPE samples selectively impairs the detection of mitochondrial inner membrane proteins and other hydrophobic complexes.

## ASSOCIATED CONTENT

### Data availability

The mass spectrometry proteomics data related to this study have been deposited to iProX^38^ database (https://www.iprox.cn/page/PDV0141.html) under the accession number IPX0013448000.

### Supporting Information

The Supporting Information is available free of charge at …

Partial Experimental section, partial results (Figures S1−S5) of method evaluation and classifier performance comparison (PDF).

### Author Contributions

Chuping Wei: Methodology, Investigation, Formal analysis. Qiuxia Zhang: Methodology, Investigation. Changying Fu: Investigation. Yeye Leng: Investigation. Chuanxi Huang: Formal analysis. Fuchu He: Supervision. Yun Yang: Conceptualization, Supervision, Writing – Review & Editing.

## ACKNOWLEDGMENTS

This work was supported by the Ministry of Science and Technology of the People’s Republic of China (Grant No. 2020YFE0202200), the Pre-study Project of Phronesis Medicine Large-scale Scientific Facility funded by Guangzhou Development District, and the National Natural Science Foundation of China (Grant No. 32501326).

## Notes

The authors declare no competing financial interest.

## For TOC

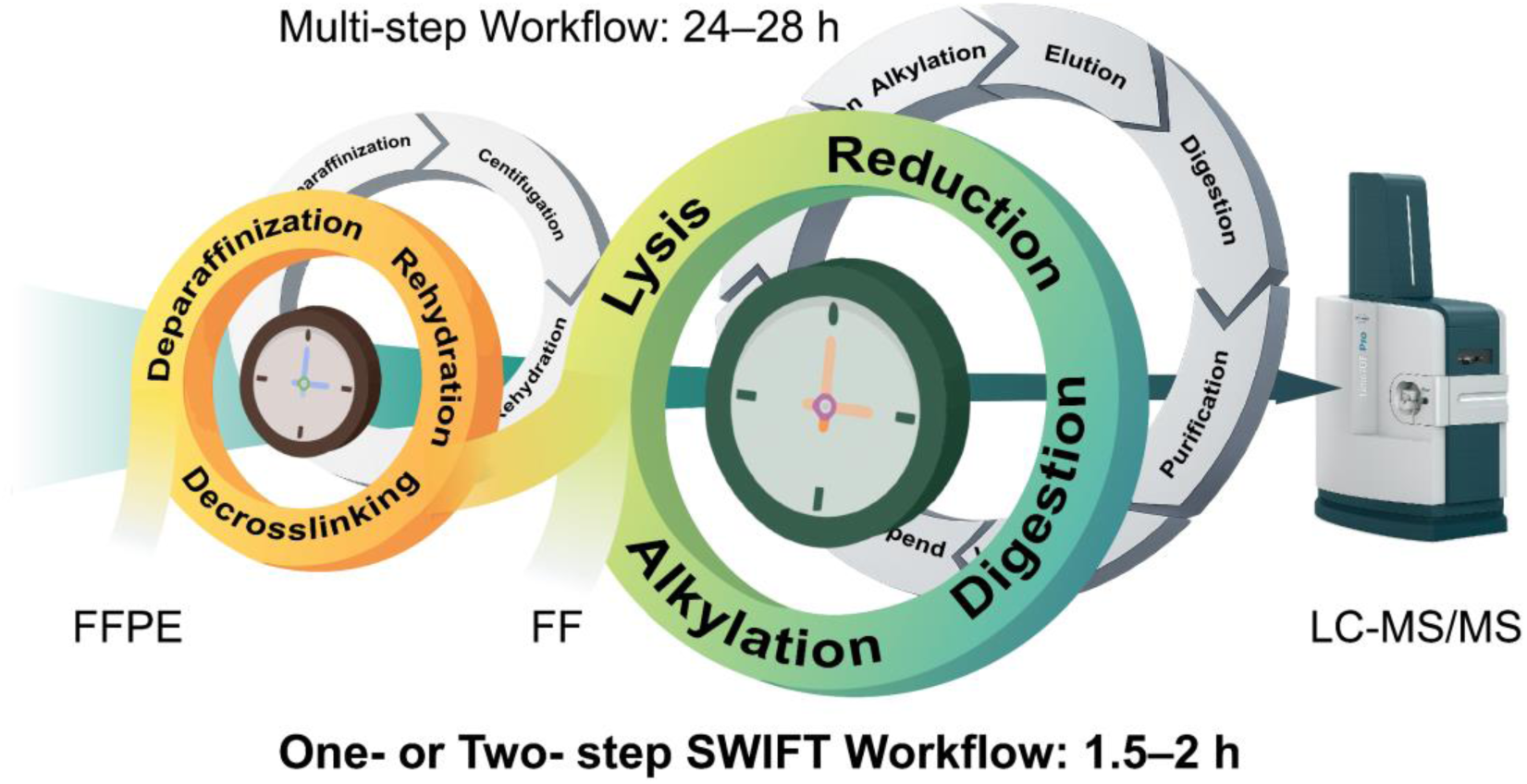

## REFERENCES

(1) Guo, T.; Steen, J. A.; Mann, M. Mass-Spectrometry-Based Proteomics: From Single Cells to Clinical Applications. Nature 2025, 638 (8052), 901–911. 10.1038/s41586-025-08584-0.

(2) Aebersold, R.; Mann, M. Mass Spectrometry-Based Proteomics. 2003.

(3) Ong, S.-E.; Mann, M. Mass Spectrometry–Based Proteomics Turns Quantitative. Nat. Chem. Biol. 2005, 1 (5), 252–262. 10.1038/nchembio736.

(4) Zhang, Y.; Fonslow, B. R.; Shan, B.; Baek, M.-C.; Yates, J. R. Protein Analysis by Shotgun/Bottom-up Proteomics. Chem. Rev. 2013, 113 (4), 2343–2394. 10.1021/cr3003533.

(5) Hughes, C. S.; Foehr, S.; Garfield, D. A.; Furlong, E. E.; Steinmetz, L. M.; Krijgsveld, J. Ultrasensitive Proteome Analysis Using Paramagnetic Bead Technology. Mol. Syst. Biol. 2014, 10 (10), 757. 10.15252/msb.20145625.

(6) Wiśniewski, J. R.; Zougman, A.; Nagaraj, N.; Mann, M. Universal Sample Preparation Method for Proteome Analysis. Nat. Methods 2009, 6 (5), 359–362. 10.1038/nmeth.1322.

(7) Kulak, N. A.; Pichler, G.; Paron, I.; Nagaraj, N.; Mann, M. Minimal, Encapsulated Proteomic-Sample Processing Applied to Copy-Number Estimation in Eukaryotic Cells. Nat. Methods 2014, 11 (3), 319–324. 10.1038/nmeth.2834.

(8) Chen, W.; Wang, S.; Adhikari, S.; Deng, Z.; Wang, L.; Chen, L.; Ke, M.; Yang, P.; Tian, R. Simple and Integrated Spintip-Based Technology Applied for Deep Proteome Profiling. Anal. Chem. 2016, 88 (9), 4864–4871. 10.1021/acs.analchem.6b00631.

(9) Shao, S.; Guo, T.; Gross, V.; Lazarev, A.; Koh, C. C.; Gillessen, S.; Joerger, M.; Jochum, W.; Aebersold, R. Reproducible Tissue Homogenization and Protein Extraction for Quantitative Proteomics Using MicroPestle-Assisted Pressure-Cycling Technology. J. Proteome Res. 2016, 15 (6), 1821–1829. 10.1021/acs.jproteome.5b01136.

(10) Xavier, D.; Lucas, N.; Williams, S. G.; Koh, J. M. S.; Ashman, K.; Loudon, C.; Reddel, R.; Hains, P. G.; Robinson, P. J. Heat ‘n Beat: A Universal High-Throughput End-to-End Proteomics Sample Processing Platform in under an Hour. Anal. Chem. 2024, 96 (10), 4093–4102. 10.1021/acs.analchem.3c04708.

(11) Lu, T.; Qian, L.; Xie, Y.; Zhang, Q.; Liu, W.; Ge, W.; Zhu, Y.; Ma, L.; Zhang, C.; Guo, T. Tissue-Characteristic Expression of Mouse Proteome. Mol. Cell. Proteomics 2022, 21 (10), 100408. 10.1016/j.mcpro.2022.100408.

12. (12) Doellinger, J. Sample Preparation by Easy Extraction and Digestion (SPEED) - A Universal, Rapid, and Detergent-Free Protocol for Proteomics Based on Acid Extraction*.

(13) Jiang, X.; Jiang, X.; Feng, S.; Tian, R.; Ye, M.; Zou, H. Development of Efficient Protein Extraction Methods for Shotgun Proteome Analysis of Formalin-Fixed Tissues. J. Proteome Res. 2007, 6 (3), 1038–1047. 10.1021/pr0605318.

(14) Ostasiewicz, P.; Zielinska, D. F.; Mann, M.; Wiśniewski, J. R. Proteome, Phosphoproteome, and N-Glycoproteome Are Quantitatively Preserved in Formalin-Fixed Paraffin-Embedded Tissue and Analyzable by High-Resolution Mass Spectrometry. J. Proteome Res. 2010, 9 (7), 3688–3700. 10.1021/pr100234w.

(15) Zhu, Y.; Weiss, T.; Zhang, Q.; Sun, R.; Wang, B.; Yi, X.; Wu, Z.; Gao, H.; Cai, X.; Ruan, G.; Zhu, T.; Xu, C.; Lou, S.; Yu, X.; Gillet, L.; Blattmann, P.; Saba, K.; Fankhauser, C. D.; Schmid, M. B.; Rutishauser, D.; Ljubicic, J.; Christiansen, A.; Fritz, C.; Rupp, N. J.; Poyet, C.; Rushing, E.; Weller, M.; Roth, P.; Haralambieva, E.; Hofer, S.; Chen, C.; Jochum, W.; Gao, X.; Teng, X.; Chen, L.; Zhong, Q.; Wild, P. J.; Aebersold, R.; Guo, T. High-throughput Proteomic Analysis of FFPE Tissue Samples Facilitates Tumor Stratification. Mol. Oncol. 2019, 13 (11), 2305–2328. 10.1002/1878-0261.12570.

(16) Mitsa, G.; Guo, Q.; Goncalves, C.; Preston, S. E. J.; Lacasse, V.; Aguilar-Mahecha, A.; Benlimame, N.; Basik, M.; Spatz, A.; Batist, G.; Miller, W. H.; Del Rincon, S. V.; Zahedi, R. P.; Borchers, C. H. A Non-Hazardous Deparaffinization Protocol Enables Quantitative Proteomics of Core Needle Biopsy-Sized Formalin-Fixed and Paraffin-Embedded (FFPE) Tissue Specimens. Int. J. Mol. Sci. 2022, 23 (8), 4443. 10.3390/ijms23084443.

(17) Kuras, M.; Woldmar, N.; Kim, Y.; Hefner, M.; Malm, J.; Moldvay, J.; Döme, B.; Fillinger, J.; Pizzatti, L.; Gil, J.; Marko-Varga, G.; Rezeli, M. Proteomic Workflows for High-Quality Quantitative Proteome and Post-Translational Modification Analysis of Clinically Relevant Samples from Formalin-Fixed Paraffin-Embedded Archives. J. Proteome Res. 2021, 20 (1), 1027–1039. 10.1021/acs.jproteome.0c00850.

(18) Marchione, D. M.; Ilieva, I.; Devins, K.; Sharpe, D.; Pappin, D. J.; Garcia, B. A.; Wilson, J. P.; Wojcik, J. B. HYPERsol: High-Quality Data from Archival FFPE Tissue for Clinical Proteomics. J. Proteome Res. 2020, 19 (2), 973–983. 10.1021/acs.jproteome.9b00686.

19. (19) Török, B.; Schäfer, C.; Kokel, A. Chapter 3.7 - Multicomponent Reactions. In Heterogeneous Catalysis in Sustainable Synthesis; Török, B., Schäfer, C., Kokel, A., Eds.; Elsevier, 2022; pp 443–489. 10.1016/B978-0-12-817825-6.00002-1.

(20) Patel, P.; Vaghani, H.; Kumbhani, J.; Kardani, H.; Patel, S.; Patel, Sh. Efficiency and Diversity in Chemical Synthesis: Exploring One-Pot Multicomponent Reactions. Russ. J. Org. Chem. 2024, 60 (9), 1752–1760. 10.1134/S1070428024090185.

(21) Uhlén, M.; Fagerberg, L.; Hallström, B. M.; Lindskog, C.; Oksvold, P.; Mardinoglu, A.; Sivertsson, Å.; Kampf, C.; Sjöstedt, E.; Asplund, A.; Olsson, I.; Edlund, K.; Lundberg, E.; Navani, S.; Szigyarto, C. A.-K.; Odeberg, J.; Djureinovic, D.; Takanen, J. O.; Hober, S.; Alm, T.; Edqvist, P.-H.; Berling, H.; Tegel, H.; Mulder, J.; Rockberg, J.; Nilsson, P.; Schwenk, J. M.; Hamsten, M.; Von Feilitzen, K.; Forsberg, M.; Persson, L.; Johansson, F.; Zwahlen, M.; Von Heijne, G.; Nielsen, J.; Pontén, F. Tissue-Based Map of the Human Proteome. Science 2015, 347 (6220), 1260419. 10.1126/science.1260419.

(22) Gustafsson, O. J. R.; Arentz, G.; Hoffmann, P. Proteomic Developments in the Analysis of Formalin-Fixed Tissue. Biochim. Biophys. Acta BBA - Proteins Proteomics 2015, 1854 (6), 559–580. 10.1016/j.bbapap.2014.10.003.

(23) García-Vence, M.; Chantada-Vazquez, M. D. P.; Sosa-Fajardo, A.; Agra, R.; Barcia De La Iglesia, A.; Otero-Glez, A.; García-González, M.; Cameselle-Teijeiro, J. M.; Nuñez, C.; Bravo, J. J.; Bravo, S. B. Protein Extraction From FFPE Kidney Tissue Samples: A Review of the Literature and Characterization of Techniques. Front. Med. 2021, 8, 657313. 10.3389/fmed.2021.657313.

(24) Fu, Z.; Yan, K.; Rosenberg, A.; Jin, Z.; Crain, B.; Athas, G.; Heide, R. S. V.; Howard, T.; Everett, A. D.; Herrington, D.; Van Eyk, J. E. Improved Protein Extraction and Protein Identification from Archival Formalin-fixed Paraffin-embedded Human Aortas. PROTEOMICS – Clin. Appl. 2013, 7 (3–4), 217–224. 10.1002/prca.201200064.

(25) Geiszler, D. J.; Kong, A. T.; Avtonomov, D. M.; Yu, F.; Leprevost, F. D. V.; Nesvizhskii, A. I. PTM-Shepherd: Analysis and Summarization of Post-Translational and Chemical Modifications from Open Search Results. Mol. Cell. Proteomics 2021, 20, 100018. 10.1074/mcp.TIR120.002216.

(26) Chang, H.-Y.; Kong, A. T.; Da Veiga Leprevost, F.; Avtonomov, D. M.; Haynes, S. E.; Nesvizhskii, A. I. Crystal-C: A Computational Tool for Refinement of Open Search Results. J. Proteome Res. 2020, 19 (6), 2511–2515. 10.1021/acs.jproteome.0c00119.

(27) Faktor, J.; Kote, S.; Bienkowski, M.; Hupp, T. R.; Marek-Trzonkowska, N. Novel FFPE Proteomics Method Suggests Prolactin Induced Protein as Hormone Induced Cytoskeleton Remodeling Spatial Biomarker. *Commun*. Biol. 2024, 7 (1), 708. 10.1038/s42003-024-06354-8.

(28) Tayri-Wilk, T.; Slavin, M.; Zamel, J.; Blass, A.; Cohen, S.; Motzik, A.; Sun, X.; Shalev, D. E.; Ram, O.; Kalisman, N. Mass Spectrometry Reveals the Chemistry of Formaldehyde Cross-Linking in Structured Proteins. Nat. Commun. 2020, 11 (1). 10.1038/s41467-020-16935-w.

(29) Magdeldin, S.; Yamamoto, T. Toward Deciphering Proteomes of Formalin-fixed Paraffin-embedded (FFPE) Tissues. PROTEOMICS 2012, 12 (7), 1045–1058. 10.1002/pmic.201100550.

(30) Dapic, I.; Baljeu-Neuman, L.; Uwugiaren, N.; Kers, J.; Goodlett, D. R.; Corthals, G. L. Proteome Analysis of Tissues by Mass Spectrometry. Mass Spectrom. Rev. 2019, 38 (4– 5), 403–441. 10.1002/mas.21598.

(31) Sprung, R. W.; Brock, J. W. C.; Tanksley, J. P.; Li, M.; Washington, M. K.; Slebos, R. J. C.; Liebler, D. C. Equivalence of Protein Inventories Obtained from Formalin-Fixed Paraffin-Embedded and Frozen Tissue in Multidimensional Liquid Chromatography-Tandem Mass Spectrometry Shotgun Proteomic Analysis. Mol. Cell. Proteomics 2009, 8 (8), 1988–1998. 10.1074/mcp.M800518-MCP200.

32. (32) Boutet, E.; Lieberherr, D.; Tognolli, M.; Schneider, M.; Bairoch, A. UniProtKB/Swiss- Prot. In Plant Bioinformatics: Methods and Protocols; Edwards, D., Ed.; Humana Press: Totowa, NJ, 2007; pp 89–112. 10.1007/978-1-59745-535-0_4.

(33) Magdeldin, S.; Yamamoto, T. Toward Deciphering Proteomes of Formalin-Fixed Paraffin-Embedded (FFPE) Tissues. PROTEOMICS 2012, 12 (7), 1045–1058. 10.1002/pmic.201100550.

(34) Hallgren, J.; Tsirigos, K. D.; Pedersen, M. D.; Almagro Armenteros, J. J.; Marcatili, P.; Nielsen, H.; Krogh, A.; Winther, O. DeepTMHMM Predicts Alpha and Beta Transmembrane Proteins Using Deep Neural Networks. Bioinformatics April 10, 2022. 10.1101/2022.04.08.487609.

(35) Tsirigos, K. D.; Peters, C.; Shu, N.; Käll, L.; Elofsson, A. The TOPCONS Web Server for Consensus Prediction of Membrane Protein Topology and Signal Peptides. Nucleic Acids Res. 2015, 43 (W1), W401–W407. 10.1093/nar/gkv485.

(36) Bernsel, A.; Viklund, H.; Falk, J.; Lindahl, E.; Von Heijne, G.; Elofsson, A. Prediction of Membrane-Protein Topology from First Principles. Proc. Natl. Acad. Sci. 2008, 105 (20), 7177–7181. 10.1073/pnas.0711151105.

(37) Teufel, F.; Almagro Armenteros, J. J.; Johansen, A. R.; Gíslason, M. H.; Pihl, S. I.; Tsirigos, K. D.; Winther, O.; Brunak, S.; Von Heijne, G.; Nielsen, H. SignalP 6.0 Predicts All Five Types of Signal Peptides Using Protein Language Models. Nat. Biotechnol. 2022, 40 (7), 1023–1025. 10.1038/s41587-021-01156-3.

(38) Ma, J.; Chen, T.; Wu, S.; Yang, C.; Bai, M.; Shu, K.; Li, K.; Zhang, G.; Jin, Z.; He, F.; Hermjakob, H.; Zhu, Y. iProX: An Integrated Proteome Resource. Nucleic Acids Res. 2019, 47 (D1), D1211–D1217. 10.1093/nar/gky869.

