## Supplementary Material for "Simple, Fast and Highly Efficient One-or Two-step Proteomic Preparation Enables Deep Profiling of Microgram-level FF and FFPE Tissues"

**Contents**


**Chemicals and Reagents** S3

**Mouse Organ Tissue Resources** S3

**NanoLC-MS/MS analysis on timsTOF Pro** S3

**Data analysis** S4

**Bioinformatics and statistical analysis** S5

**Figure S1.** S8

**Figure S2.** S9

**Figure S3.** S10

**Figure S4.** S11

**Figure S5.** S13

**References** S14

### ■ EXPERIMENTAL SECTION

#### Chemicals and Reagents

Liquid chromatography–mass spectrometry (LC–MS) grade water and acetonitrile (ACN) were purchased from Fisher Chemical (USA). Formic acid (FA) was purchased from Fluka (UK). The Milli-Q system (Millipore, Billerica, MA, USA) was used to prepare ultrapure water. Phosphate-buffered saline (PBS) and tris(hydroxymethyl)aminomethane (Tris)–HCl buffers were purchased from Solarbio (Beijing, China). 2-chloroacetamide (CAA), sodium deoxycholate (SDC), and tris(2-carboxyethyl)phosphine hydrochloride (TCEP-HCl) were purchased from Sigma-Aldrich (St. Louis, MO, USA). n-dodecyl-β-D-maltoside (DDM) was purchased from Energy Chemical (China). Rapid trypsin/Lys-C digestion kits were purchased from Promega (Madison, WI, USA). Pierce™ HeLa Protein Digest Standard was purchased from Thermo Scientific (Waltham, MA, USA). Deparaffinization Solution was purchased from Baso (Zhuhai, China).

#### Mouse Organ Tissue Resources

All animal procedures were approved by the Institutional Animal Care and Use Committee (IACUC) of the Southern University of Science and Technology. Whole organs were collected from adult female C57BL/6J mouse after perfusion with normal saline (0.9% NaCl) to minimize blood-protein contamination. Brain, heart, spleen, lung, kidney, liver, and pancreas were harvested. Fresh frozen (FF) specimens were snap frozen in liquid nitrogen and stored at −80 °C. Formalin-fixed, paraffin-embedded (FFPE) specimens were immersed in 10% neutral buffered formalin (NBF) and fixed for ~16–24 h at 22 °C, then processed by graded-ethanol dehydration, xylene clearing, paraffin infiltration, and embedding. For FF method development, liver tissues were used; for FFPE method development, brain tissues were used. For the direct comparison between preservation methods, each organ was bisected along its largest cross-sectional plane: one half was embedded in optimal cutting temperature (OCT) compound and snap-frozen in liquid nitrogen, and the other half was formalin-fixed and paraffin-embedded. Sections were cut at 10 μm for FF tissues and 4 μm for FFPE tissues.

#### LC-MS/MS analysis on timsTOF Pro

Nano liquid chromatography-tandem mass spectrometry (nanoLC-MS/MS) analysis was conducted employing a hybrid trapped ion mobility spectrometry (TIMS) quadrupole time-of-flight mass spectrometer (timsTOF Pro, Bruker Daltonics) along with a CaptiveSpray nano-electrospray ion source. Peptide samples underwent separation using an UltiMate 3000 RSLCnano system (Thermo Scientific) connected online to the timsTOF Pro. This system employed integrated spraytip columns (100 μm i.d. x 30 cm) packed with 1.9 μm/120 Å ReproSil-Pur C18 resins (Dr. Maisch GmbH, Germany). Peptide separation was accomplished over a 140-min total gradient for method development of One-step SWIFT for FF tissues: 0%–4% (v/v) buffer B over 2 min, 4%–25% (v/v) buffer B over 99 min, 25%–40% (v/v) buffer B over 20 min, 40%–99% (v/v) buffer B over 10 min, and 99% (v/v) buffer B was held for 10 min. The analytical flow rate was 0.3 μL/min except that 0.6 μL /min was applied in the first 2 min. For method development of Two-step SWIFT for FFPE tissues, and multi-organ proteomics analysis, a 110-min total gradient was applied as follows: 2.2%-4.5% (v/v) buffer B over 5 min, 4.5% -24.7% (v/v) buffer B over 69 min, 24.7% -39.3% (v/v) buffer B over 15 min, 39.3% -99% (v/v) buffer B over 10 min, and 99% (v/v) buffer B was held for 10 min. The analytical flow rate was 0.3 μL/min except that 0.6 μL /min was applied in the first 5 min.

Mass spectrometry analysis was performed in positive-ion data-independent acquisition parallel accumulation–serial fragmentation (diaPASEF) mode with TIMS enabled, scanning precursors across an m/z range of 300–1,500. MS2 spectra were acquired with m/z 400 to 1200, with quadrupole isolation width 25 Th using a scheme of 64 windows (16 scans per MS1, 4 steps per scan), yielding a total cycle time of 1.8 s. The ion-mobility range was set to $1/K_0$ = 1.40 to 0.75 V·s·cm$^{-2}$, and the ramp time was 100 ms. During diaPASEF MS/MS acquisition, the collision energy was fixed at 10 eV. Source parameters were as follows: capillary 4.5 kV, end-plate offset 500 V, nebulizer 0.4 bar, dry gas 3.0 L·min$^{-1}$, and dry temperature 180 °C.

For data dependent acquisition (DDA)-MS analyses, the elution gradient was set to: 0%-4% (v/v) buffer B over 2 min, 4% -25% (v/v) buffer B over 74 min, 25% -40% (v/v) buffer B over 15 min, 40% -99% (v/v) buffer B over 10 min, and 99% (v/v) buffer B was held for 10 min. Columns were heated at 60°C. The flow rate during gradient elution was maintained at 0.3 μL/min, but was increased to 0.6 μL /min for the initial 2 min. MS1 spectra were acquired with m/z 300–1,500; ion mobility ($1/K_0$) range was 0.75–1.30 V·s cm$^{-2}$; ramp time was 166 ms. Charge-state range was set to 2–5; 4

PASEF scans were followed to each MS1 scan. Target intensity of 20,000 (arbitrary units) and intensity threshold of 2,500 were used, giving a total cycle time of 0.86 s.

**Data analysis**

DIA data from method development were analyzed using Spectronaut 19.4 (Biognosys) software in directDIA mode. These data were searched against with Swiss-Prot database (17,090 entries, downloaded on December 3, 2021, for mouse tissue samples and 20,375 entries, downloaded on December 3, 2021 for commercial Hela digest samples). Cysteine carbamidomethylation was set as fixed modification. N-terminal acetylation of protein and methionine oxidation were set as variable modifications. Two missed cleavages were allowed with trypsin digestion. Precursor filtering was set to perform based on Q-values. Quantification results were output with protein and peptide false discovery rates (FDRs) less than 0.01. All other settings were left at their default values. Alkylation reaction efficiency was evaluated by setting +57 Da carbamidomethylation as a variable modification and calculating the percentage of Cys-containing peptides that carried this modification.

DDA data for formaldehyde-induced modifications in FFPE samples were profiled by open search using MSFragger (v3.8) via FragPipe (v20) with default parameters. PeptideProphet was used for PSM validation, and PTM-Shepherd (default, in FragPipe) performed mass-shift profiling and annotation.

**Bioinformatics and statistical analysis**

For multi-organ proteomics benchmarking of the SWIFT workflows, FF and FFPE tissue datasets were analyzed independently. Protein abundance values were $\log_2$-transformed, median-centered and global-minimum imputation. Within each organ, quantified proteins not present in over half samples were filtered. Principal component analysis was performed using probabilistic PCA (PPCA) as implemented in the pcaMethods R package[1]; variance explained was derived via explVar. Coefficients of variation (CV) were calculated but only reported when at least three replicates per organ were available.

Differential expression was assessed using limma[2] with empirical Bayes variance moderation and TREAT tests against predefined fold-change thresholds. For each organ, pairwise contrasts were performed against all others ($|\log_2 FC| \geq 1$, trend=TRUE). GO:BP enrichment was conducted on significant proteins, and terms were ranked by Benjamini–Hochberg global FDR (q global) [3]; the top 10 terms per organ were retained, with intersection size used to resolve ties.

For FF vs. FFPE comparisons, the same preprocessing workflow was applied, with an additional 2/3 presence filter. Hypothesis testing used TREAT ($|\log_2 FC| \geq 0.8$, trend=TRUE) for global FF or FFPE enriched proteins. Proteins were classified as FF-enriched ($\log_2 FC > 0$) or FFPE-enriched ($\log_2 FC < 0$). Significance was controlled at FDR $\leq 0.05$.

Globally FF-enriched and FFPE-enriched UniProt accessions were mapped to reviewed mouse (Swiss-Prot) sequences, prioritizing exact IDs (isoform-suffix fallback) and retaining the longest sequence[4]. For each sequence we computed isoelectric point (pI), GRAVY (mean Kyte–Doolittle hydropathy), and lysine content (%)[5]. Group differences (FF-enriched vs FFPE-enriched) were assessed per metric by the two-sided Wilcoxon rank-sum test, and effect size reported as Cliff's delta. Proteins were annotated with SignalP 6.0[6] for N-terminal signal peptides and with DeepTMHMM[7], SCAMPI[8], and TOPCONS[9] for transmembrane (TM) topology. UniProt accessions were normalized by removing isoform suffixes before merging predictions. A consensus classification was assigned by majority vote across the TM predictors, combined with the SignalP call: proteins were labeled as Membrane (TM only), Secreted (SP only), SP+TM (both), or Other (neither).

Within each organ, proteins were considered preservation method-enriched when significantly up-regulated with $\log_2 FC > 1.0$ at FDR $\leqslant 0.01$ (Benjamini–Hochberg). Enrichment was performed using g:Profiler (hypergeometric test)[10]. Terms were retained if annotated to 10–200 genes, with ≥3 hits and background coverage ≤10% (BP/Reactome ≤500 genes; CC ≤1,000 genes). Redundant terms were collapsed to representative sets. For cross-organ summaries, P values were combined using Fisher's method, and top-ranked terms were reported.

Data analysis and visualization were performed using custom R (v4.5.1) scripts. The following R packages were used: pheatmap (v1.0.13), limma (v3.64.3), ggplot2 (v4.0.0), org.Mm.eg.db (v3.21.0), AnnotationDbi (v1.70.0), ggrepel (v0.9.6), pcaMethods (v2.0.0), tidyverse (v2.0.0), and stringr (v1.5.2). For GO analysis, g:Profiler (g:GOSt) was run for *Mus musculus* with Benjamini–Hochberg FDR control ($q < 0.05$). Statistical analyses and plots for the five reported markers were created using GraphPad Prism (v10).

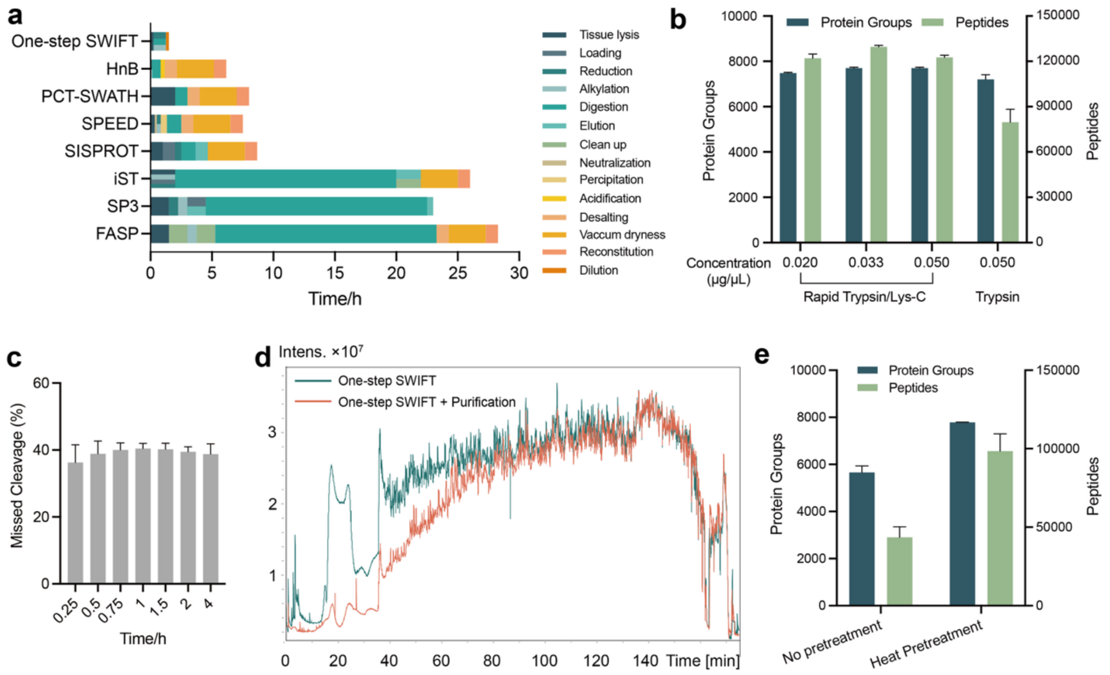


**Figure S1. Evaluation and benchmarking of the One-step SWIFT workflow.** (a) Comparison of processing time and step count for tissue proteomic preparation (20–40 samples in a batch). (b) Missed cleavage rates across digestion times (0.25–4 h). (c) Total ion chromatograms (TICs) for One-step SWIFT and One-step SWIFT with extra purification. (e) Effect of pretreatment on pancreas tissues: high-temperature pretreatment increased the number of protein group and peptide identifications.

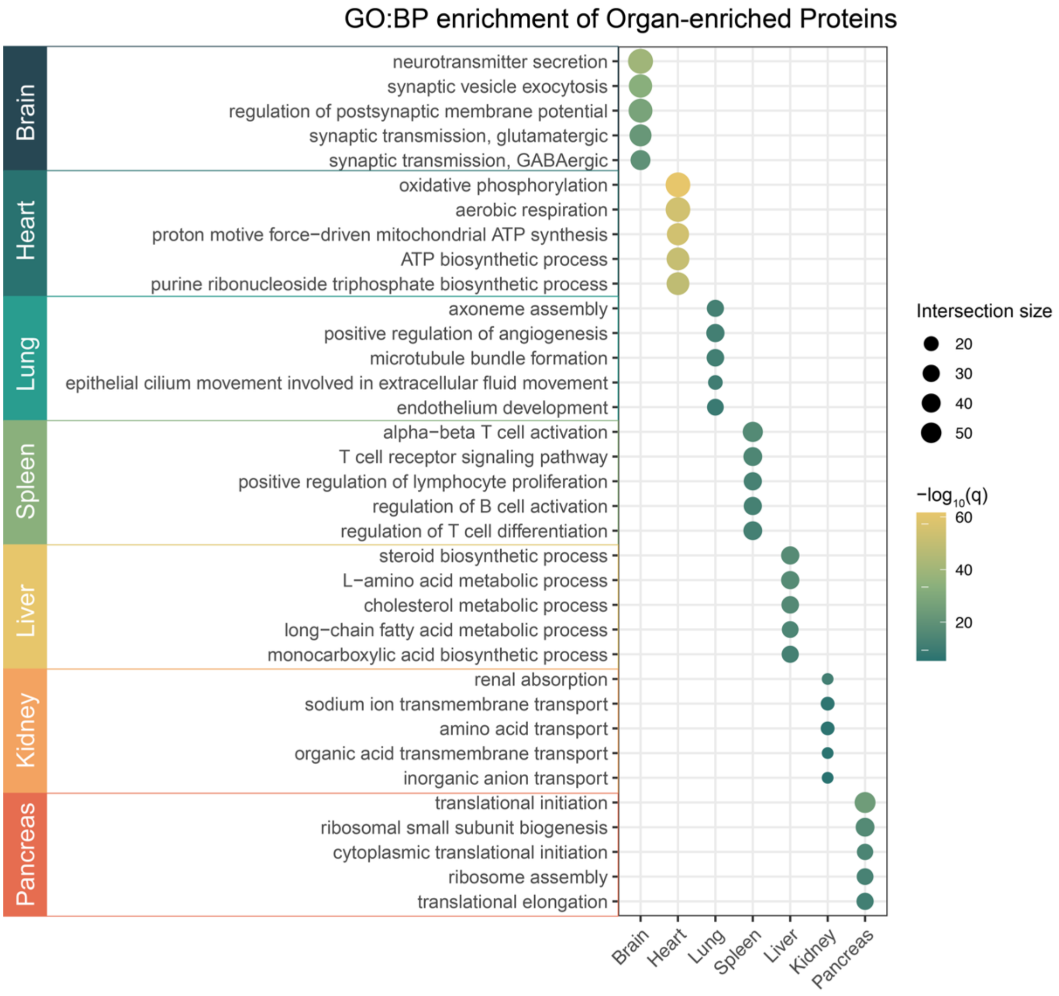


**Figure S2.** GO biological process enrichment of organ-specific protein sets. Redundant terms were removed; the top representative terms are shown. Dot color indicates $-\log_{10}(q)$ and dot size represents the intersection size.

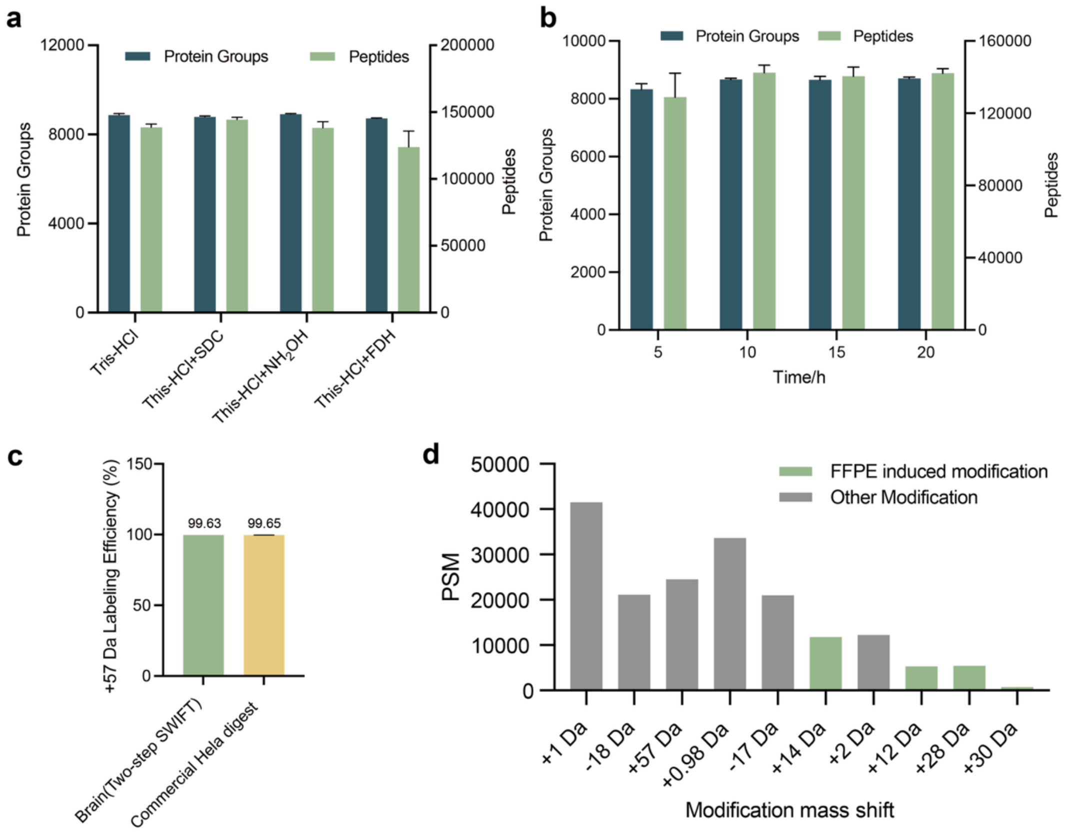


**Figure S3. Development and evaluation of the Two-step SWIFT workflow.** (a) Protein group and peptide identifications with SDC, hydroxylamine, or FDH additives. (b) Protein group and peptide identifications across different incubation times. (c) Cysteine alkylation (+57 Da) efficiency in FFPE brain samples prepared by Two-step-SWIFT versus a commercial HeLa digest. (d) The ten most abundant mass shifts in FFPE brain tissues, shown as percentages of total PSMs; Fixation-related modifications are labelled in green.

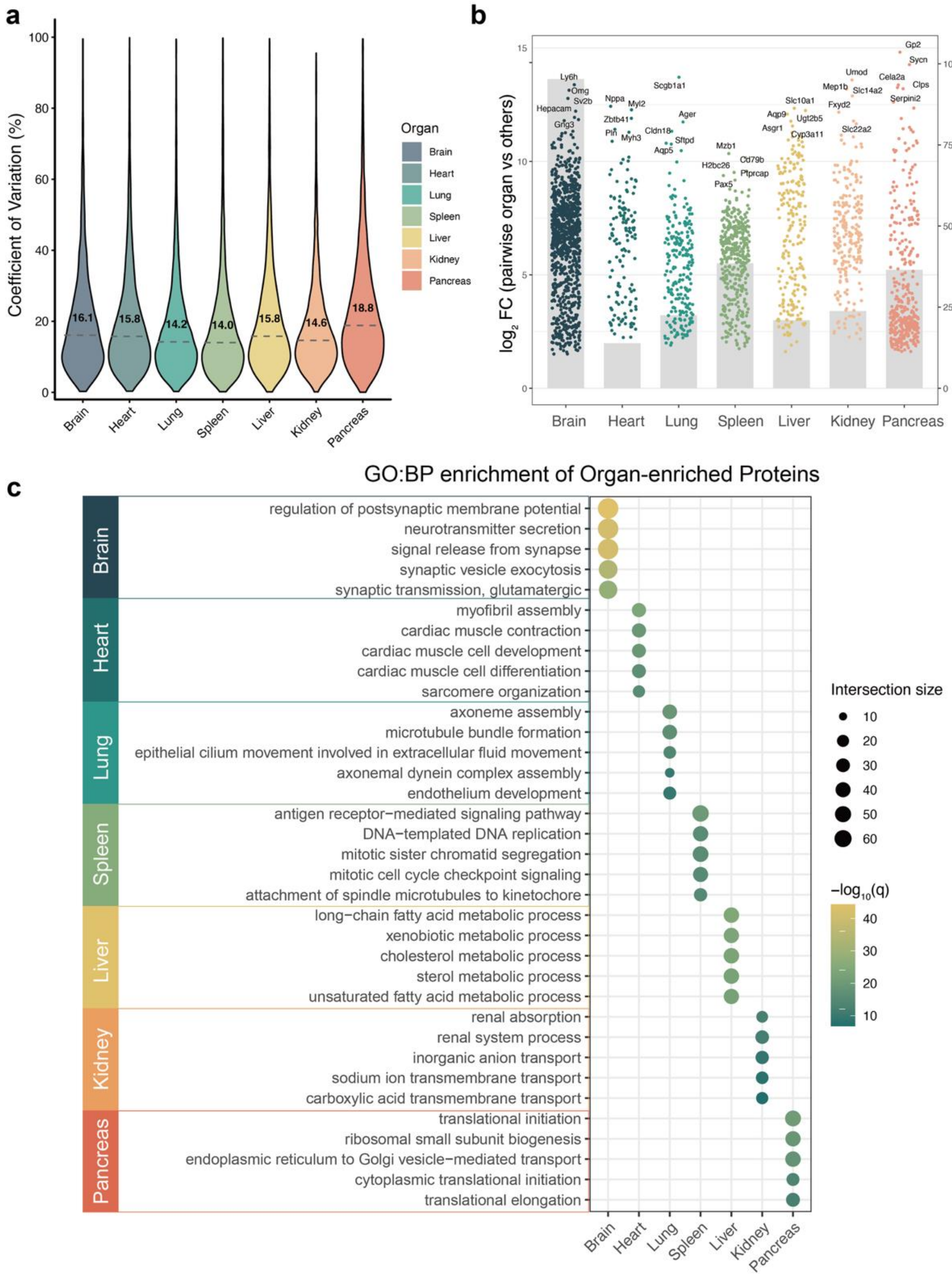


**Figure S4. Comprehensive benchmarking of FFPE multi-organ proteomics analysis.** (a) Coefficient of variation (CV) percentages across organ using the conventional multi-step workflow; medians are indicated in violin plots. (b) Volcano plots of pairwise organ-specific proteins (FDR $\leqslant 0.01$, $|\log_2 FC| \geqslant 1$) for FFPE tissues. The top five proteins are labelled; organ-specific set sizes (protein counts) are shown on the plots. (c) GO biological process enrichment of organ-specific protein sets.

Redundant terms were removed and the top representative terms are shown. Dot color indicates $–\log_{10}(q)$ and dot size represents the intersection size.

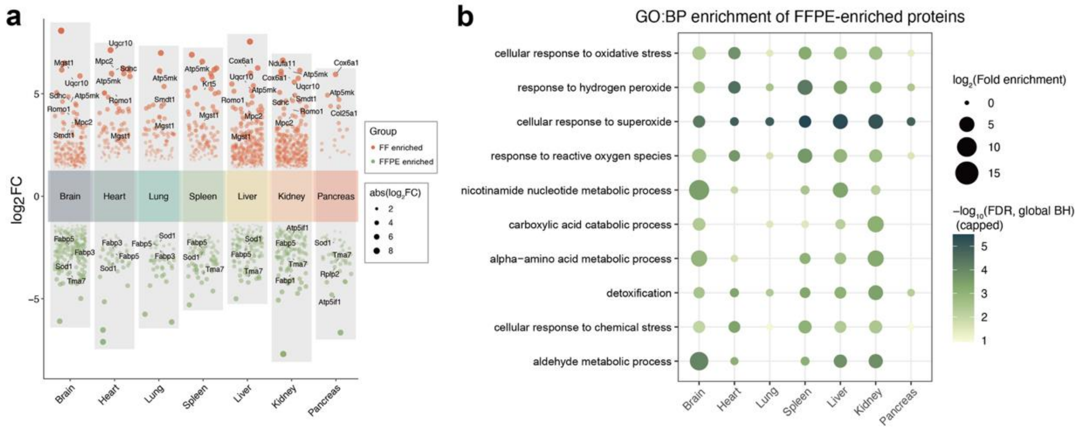


**Figure S5**. **Fixation-driven proteome differences across organs.** (a) Multi- organ differential abundance analysis ($|\log_2 FC| \geqslant 1.0$, FDR $\leqslant 0.01$) using limma; positive $\log_2 FC$ denotes FF-enriched and negative denotes FFPE-enriched proteins. (b) GO biological process over-representation analysis for FF-enriched proteins by organ.

### REFERENCES


(1) Stacklies, W.; Redestig, H.; Scholz, M.; Walther, D.; Selbig, J. pcaMethods—a Bioconductor Package Providing PCA Methods for Incomplete Data. *Bioinformatics* **2007**, *23* (9), 1164–1167. https://doi.org/10.1093/bioinformatics/btm069.

(2) McCarthy, D. J.; Smyth, G. K. Testing Significance Relative to a Fold-Change Threshold Is a TREAT. *Bioinformatics* **2009**, *25* (6), 765–771. https://doi.org/10.1093/bioinformatics/btp053.

(3) Benjamini, Y.; Hochberg, Y. Controlling the False Discovery Rate: A Practical and Powerful Approach to Multiple Testing. *J. R. Stat. Soc. Ser. B Methodol.* **1995**, *57* (1), 289–300. https://doi.org/10.1111/j.2517-6161.1995.tb02031.x.

(4) Boutet, E.; Lieberherr, D.; Tognolli, M.; Schneider, M.; Bairoch, A. UniProtKB/Swiss-Prot. In *Plant Bioinformatics: Methods and Protocols*; Edwards, D., Ed.; Humana Press: Totowa, NJ, 2007; pp 89–112. https://doi.org/10.1007/978-1-59745-535-0_4.

(5) Kyte, J.; Doolittle, R. F. A Simple Method for Displaying the Hydropathic Character of a Protein. *J. Mol. Biol.* **1982**, *157* (1), 105–132. https://doi.org/10.1016/0022-2836(82)90515-0.

(6) Teufel, F.; Almagro Armenteros, J. J.; Johansen, A. R.; Gíslason, M. H.; Pihl, S. I.; Tsirigos, K. D.; Winther, O.; Brunak, S.; Von Heijne, G.; Nielsen, H. SignalP 6.0 Predicts All Five Types of Signal Peptides Using Protein Language Models. *Nat. Biotechnol.* **2022**, *40* (7), 1023–1025. https://doi.org/10.1038/s41587-021-01156-3.

(7) Hallgren, J.; Tsirigos, K. D.; Pedersen, M. D.; Almagro Armenteros, J. J.; Marcatili, P.; Nielsen, H.; Krogh, A.; Winther, O. DeepTMHMM Predicts Alpha and Beta Transmembrane Proteins Using Deep Neural Networks. Bioinformatics April 10, 2022. https://doi.org/10.1101/2022.04.08.487609.

(8) Bernsel, A.; Viklund, H.; Falk, J.; Lindahl, E.; Von Heijne, G.; Elofsson, A. Prediction of Membrane-Protein Topology from First Principles. *Proc. Natl. Acad. Sci.* **2008**, *105* (20), 7177–7181. https://doi.org/10.1073/pnas.0711151105.

(9) Tsirigos, K. D.; Peters, C.; Shu, N.; Käll, L.; Elofsson, A. The TOPCONS Web Server for Consensus Prediction of Membrane Protein Topology and Signal Peptides. *Nucleic Acids Res.* **2015**, *43* (W1), W401–W407. https://doi.org/10.1093/nar/gkv485.

(10) Raudvere, U.; Kolberg, L.; Kuzmin, I.; Arak, T.; Adler, P.; Peterson, H.; Vilo, J. G:Profiler: A Web Server for Functional Enrichment Analysis and Conversions of Gene Lists (2019 Update). *Nucleic Acids Res.* **2019**, *47* (W1), W191–W198. https://doi.org/10.1093/nar/gkz369.
